# The influence of downstream structured elements within mRNA on the dynamics of intersubunit rotation in ribosomes

**DOI:** 10.1101/2024.10.17.618931

**Authors:** Bassem Shebl, Anna Pavlova, Preston Kellenberger, Dongmei Yu, Drew E. Menke, James C. Gumbart, Peter V. Cornish

## Abstract

Proper codon/anti-codon pairing within the ribosome necessitates linearity of the transcript. Any structures formed within a messenger RNA (mRNA) must be unwound before the respective codon is interpreted. Linearity, however, is not always the norm; some intricate structures within mRNA are able to exert unique ribosome/mRNA interactions to regulate translation. Intrinsic kinetic and thermal stability in many of these structures are efficient in slowing translation causing pausing of the ribosome. Altered translation kinetics arising from atypical interactions have been shown to affect intersubunit rotation. Here, we employ single-molecule Förster Resonance Energy Transfer (smFRET), to observe changes in intersubunit rotation of the ribosome as it approaches downstream structured nucleic acid. The emergence of the hyper-rotated state is critically dependent on the distance between downstream structure and the ribosome suggesting interactions with the helicase center are allosterically coupled to intersubunit rotation. Further, molecular dynamics (MD) simulations were performed to determine ribosomal protein/mRNA interactions that may play a pivotal role in helicase activity and ultimately unwinding of downstream structure.

## Introduction

The ribosome is a ubiquitous macromolecular complex that translates the information encoded in the mRNA into functional proteins. Despite the high processivity of the ribosome, protein synthesis is not a continuous process. Intermittent pausing of the ribosome slows down or even completely halts translation is an example of discontinuity ^1–3^. Ribosomal pausing has been functionally linked to co-translational protein folding ^4^, membrane localization ^5^, modulating protein expression levels ^6^ and programmed ribosomal frameshifting (PRF) ^7,8^. A common feature in these is the utilization of a secondary RNA structure within the mRNA coding region to pause the moving ribosome, permitting cells to regulate protein production. Thus, folded secondary structures within the mRNA coding region present a thermodynamic and kinetic barrier to translation, and must be disrupted for translation to proceed.

The average diameter of the mRNA entrance tunnel (∼15 Å) is smaller than the diameter of a double stranded helix (∼ 20 Å) ^9^. Thus, for translation to proceed forward, any nucleic acid structure requires unfolding. Unwinding nucleic acid duplexes is typically carried out via dedicated helicases ^10^. In the bacterial ribosome, however, the required helicase activity is intrinsic to the macromolecular complex itself ^9^. Findings by Takyar et al., predicted the position of the helicase center of the ribosome to be within the downstream mRNA entrance tunnel, ∼11 nucleotides away from the P-site ^9^. The predicted distance, interestingly, correspond to the optimal distance for a structured RNA to be capable of inducing -1 PRF, along with the upstream slippery site ^11^. Ribosomal proteins S3, S4 and S5 encircle and interact with the incoming mRNA at the 30S entrance tunnel ^12,13^. The three proteins surrounding the incoming mRNA, resemble the well-known sliding clamps of DNA and RNA polymerases, suggestive of a similar function to S3, S4 and S5 ^14^. However, no known helicase motifs have been identified in these proteins. Furthermore, despite the ribosome itself being a GTPase, these proteins demonstrate no direct use of ATP or GTP ^9^. The highly dynamic nature of the ribosomal structure, along with the lack of evidence to any external energy sources required for the unwinding activity, suggests the involvement of conformational changes of the two subunits in engaging and unwinding downstream structures facing the ribosome. ^15,16^.

Structural and biophysical studies have shown that tRNA and mRNA translocation are accompanied by large-scale conformational changes ^9,17–22^. The most notable change is the thermally-driven counterclockwise rotation of the small 30S subunit relative to the vertical axis of the large 50S subunit ^15,17,23^. Thus, subunit ratcheting renders the ribosome between two states: the rotated and non-rotated states. Accompanied by the 30S subunit rotation is a process known as head swiveling. This head domain exhibits 5° rotation about an axis within itself ^24^. The region between the head and the body of the 30S subunit forms the mRNA entrance and exit tunnels on opposite sides of the subunit. The two tunnels were found to alternatively expand and contract with subunit rotation, presumably allowing the mRNA to advance through the ribosome and maintain a grip on the reading frame ^15^. The mRNA entrance tunnel with its circular clamp-like entrance has been proposed to host the intrinsic helicase center of the ribosome. Two distinctive mechanisms have been suggested to achieve this goal ^20^. First, the ribosome can bind and stabilize the open conformation of the duplex during helical fraying. In the second mechanism, the ribosome can mechanically pull apart the two strands through an interplay between the various conformational changes that accompany translation ^18,22,24^. Thus, it is possible that an intricate balance of conformational changes regulates which mechanism is used in the process of unwinding.

Various approaches have been employed to relate structural changes to functional activity during translation, varying from Cryo-electron microscopy and X-ray crystallography to fluorescence imaging. Using smFRET, we have shown previously that ribosomal subunits spontaneously fluctuate between two conformations, corresponding to the rotated and non-rotated states, when translating a unstructured mRNA ^17^. The presence of structured mRNA within the coding region was shown to induce further rotation of the small subunit when the structure is placed at the mRNA entrance tunnel. This creates a third conformation, the hyper-rotated state, alongside the two identified states; the rotated and non-rotated states (Figure 1)^18^. The presence of the hyper-rotated state is independent of the identity and amino-acylation status of the tRNA occupying the P site. Chen et al. showed a similar observation of a non-canonical rotated state, characterized by a long pause, coupled to programmed frameshifting and bypassing of non-coding regions ^19,21^. The structural constraints imposed by the dimensions of the mRNA entrance tunnel facilitates the presence of an optimal position for an RNA structure, allowing the ribosome to sense the structure and respond in the form of unwinding. As mentioned above, investigating the highly concerted movement of the ribosomal subunits could shed light on the unwinding mechanism of the helicase center of the ribosome. Given the uniqueness of the ribosomal helicase activity and the lack of an observed direct energy source for the process, subunit rotation, seems highly likely to be integral in the sensory mechanism by which ribosomes engage structured mRNA, and unwind it. Thus, a further quantitative analysis of the influence of structured mRNA sequences on the conformational states present and the population distribution is required.

**Figure 1.**
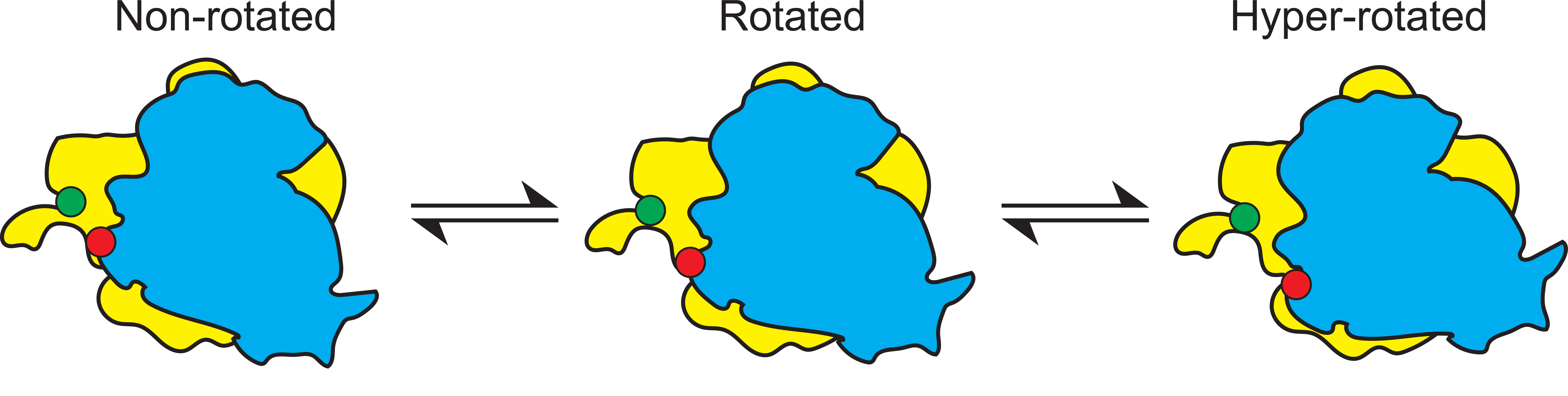
Schematic depicting the two ribosomal subunits (50S – yellow; 30S – blue) in various rotational states: non-rotated, rotated, and hyper-rotated. Approximate location of Cy3 (green sphere) attached to ribosomal protein L9 and Cy5 (red sphere) attached to ribosomal protein S6 are shown.

Here, we use a smFRET approach to track the conformational changes of intersubunit rotation during translation. Consistent with our previous work, we show that hyper-rotation is formed in the presence of secondary structures within the coding region of the mRNA at a close proximity to the ribosome ^18^. Furthermore, using a DNA-RNA hybrid duplex, we track the formation of the hyper-rotated state and illustrate the importance of the relative spatial position of a structured element on the conformational dynamics of the ribosome. We show how the presence of such a structure alters the equilibrium of the existing states towards the newly formed state; the hyper-rotated state. Alongside subunit rotation, “how does the ribosome respond near the mRNA entrance tunnel to incoming structures?”, is a standing question. Using molecular dynamics (MD) simulations, we investigated the local structural rearrangements of the clamp-like structure encircling the mRNA entrance tunnel when the ribosome encounters a structured nucleic acid at various distances. Taken together, smFRET and MD provide valuable and complementary insights into the mechanism by which the ribosome can sense incoming structured elements and how it can transduce that across the entire structure through major structural rearrangements with functional implications on translation.

## Results

### S6 (Cy5) / L9 (Cy3) ribosomes report on intersubunit rotation

For this study, ribosomal proteins S6 and L9 were individually labeled with Cy5 and Cy3, respectively, and reconstituted *in vitro* into ΔS6 / ΔL9 ribosomes purified by Ni-NTA via a 6x His tag on the L7/L12 stalk as described in Materials and Methods ^17,18,25,26^. The dual-labeled ribosomal constructs were assembled with an unstructured control mRNA employed in previous studies^9,17,25^, m291, and immobilized to the passivated surface of a quartz slide via a biotin-neutravidin linkage. Time trajectories of the individual molecules showed fluctuations between two conformations / FRET states; the non-rotated (∼ 0.58) and rotated (∼ 0.38) states as described previously (Figure S1) ^17^. The two conformations coexist in equilibrium at a ratio highly dependent on the occupancy of the A-site tRNA and the acylation status of the P-site tRNA. Ribosomes assembled with a single deacylated tRNA^fMet^ in the P site bias the conformational equilibrium towards the rotated state. Using deacylated tRNA^fMet^ provides the highest rotational freedom for the ribosome among different tested conditions as reflected in the number of dynamic molecules ^17^. In agreement with previously reported data on m291, we showed that the presence of a deacyated tRNA^fMet^ in the P site, regardless of the absence or presence of a A-site tRNA, shifted the existing equilibrium towards the rotated conformation (∼72 and 68%, respectively) ^17,26^.

### Visualization of the hyper-rotated state

Previously, we reported an additional conformational state of the 50S and 30S ribosomal subunits while interacting with the *dnaX* hairpin (Figure 1) ^18^. The *dnaX* hairpin (Figure 2C) has been well characterized and used for studying ribosome dynamics whether in isolation ^18^ or in the context of a full frameshifting signal ^19,27,28^. We observed that the placement of a structured RNA sequence within the mRNA and in close proximity to the mRNA entrance tunnel of the ribosome induces a new population with a significant reduction in FRET (0.22) as compared to the rotated (0.38) and non-rotated (0.58) states (Figure S1). Characterization of the hyper-rotated conformation showed that it has a substantial influence on the equilibrium among the different conformational states of the ribosome. Furthermore, ribosomes showed less fluctuation events in comparison to the control; indicative of a decrease in molecular dynamicity (See “The kinetics of intersubunit rotation in the presence of a folded structure”).

**Figure 2.**
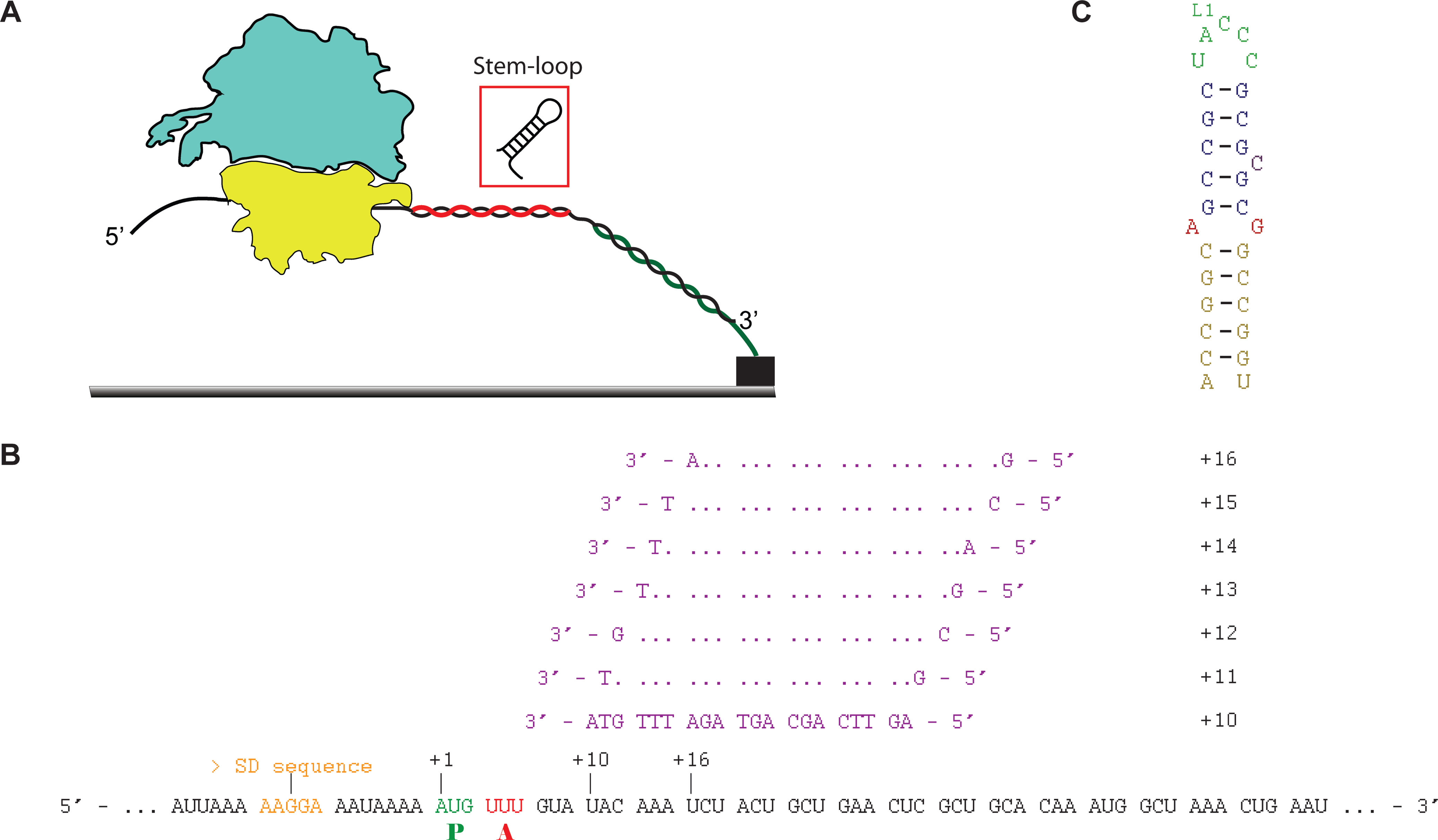
A) Ribosomes were immobilized by the hybridization of the 3’ end of the mRNA to a complementary biotinylated DNA oligo bound to the slide surface via a neutravidin-biotin-PEG linkage. Alternatively, an mRNA with *dnaX* hairpin was used. B) The sequence of m291 (mRNA) and oligonucleotides used. The nucleotide register is shown starting at the first nucleotide in the P site as +1. Oligonucleotides are labeled as +10 up to + 16 in single nucleotide increments. SD sequence, codons occupying the P and A sites are highlighted. C) The sequence of the GC-rich *dnaX* hairpin.

His-tagged, tight coupled ribosomes show more activity compared to the ribosomes employed in the past study ^31^, thus we are reproducing these experiments to provide consistency throughout the results in the present study ^18^. We assembled the His-tagged S6 (Cy5) /L9 (Cy3) 70S ribosomes with the *dnaX* hairpin and tRNA^fMet^ in the P site. The hairpin of the *dnaX* gene was placed at the mRNA entrance tunnel using a 6 nucleotide linker ^29^.

Consistent with previous results, the *dnaX* hairpin induced a significant portion of the population of ribosomes into hyper-rotation (∼ 39%, previously 68%) and rotation (∼56%, previously 20%) (Figure 3H, Table 1) coupled with suppression of the non-rotated conformation (∼ 6%, previously 20%), which shows a significant change from observed values for the linear control sequence, m291, assembled under the same conditions (0% hyper-rotated, previously 0%)(72% rotated, previously 71%)(28% non-rotated, previously 29%) ^17,18,30^. Furthermore, the percentage of fluctuating / dynamic molecules, or molecules that switch between the non-rotated and rotated conformations, decreased to ∼ 10%, an ∼ 80% reduction as compared to the linear control (∼ 52%). Thus, constructs capable of inducing hyper-rotation seem to reduce molecular dynamicity (See “The kinetics of intersubunit rotation in the presence of a folded structure”).

**Figure 3.**
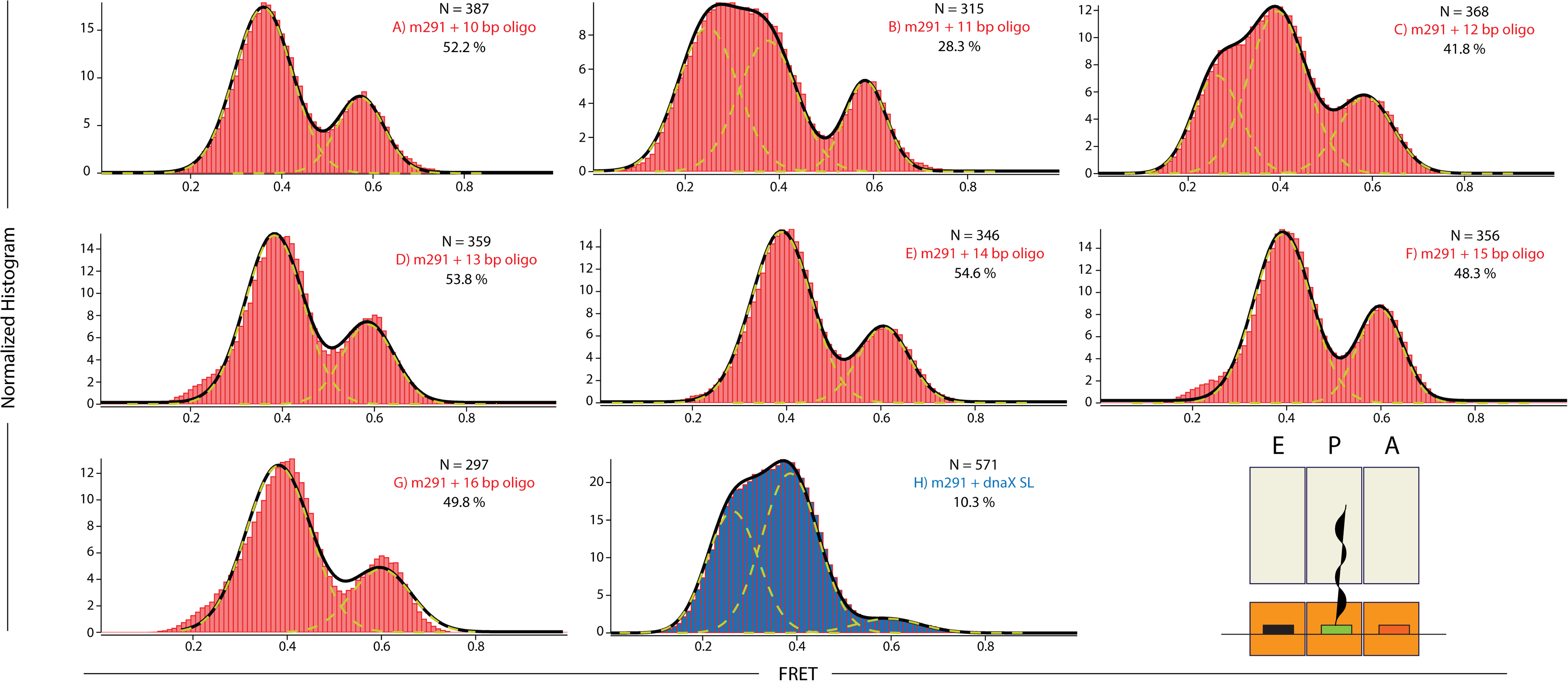
Normalized histograms complied from hundreds of traces showing distributions of FRET values for S6-Cy5/L9-Cy3 ribosomes containing a deacylated tRNA^fMet^ in the P site and nucleic acid duplexes. N number of molecules used to compile each histogram; yellow lines represent individual Gaussian fits centered at ∼ 0.26 (hyper-rotated state), 0.38 (rotated state) and 0.58 (non-rotated state) FRET efficiency, black lines represent the sum of two (A, D-G) or three (B, C and H) Gaussians, % represent the percentage of fluctuating molecules for a given ribosomal complex.

**Table 1.**
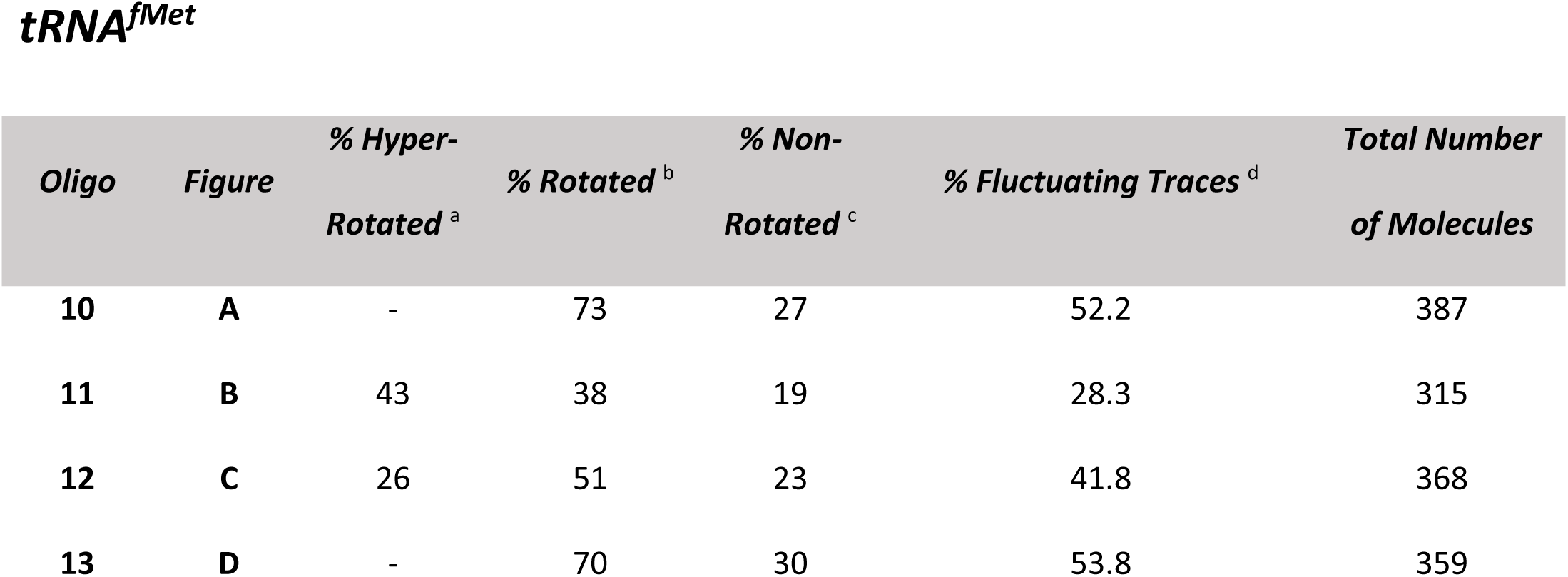
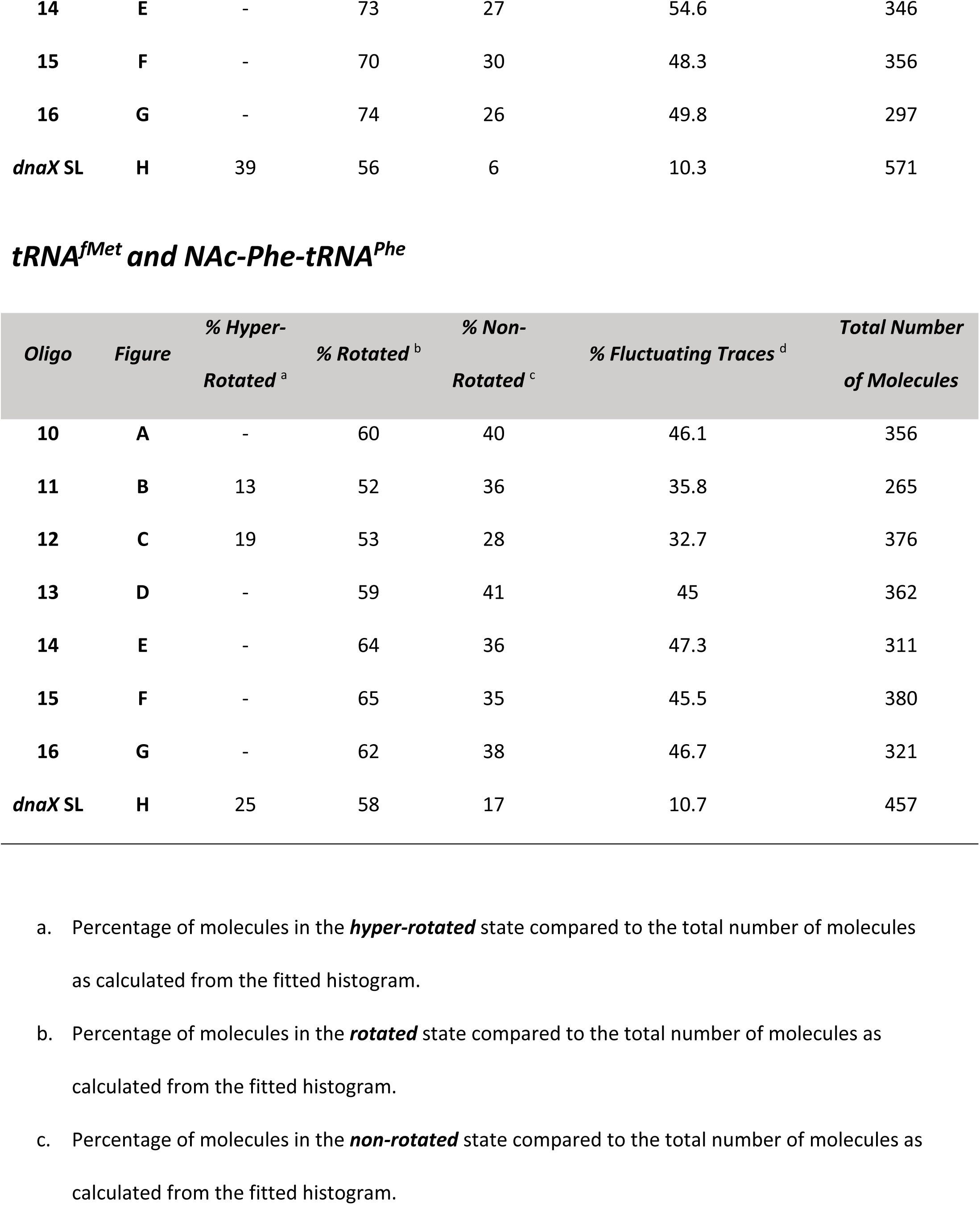
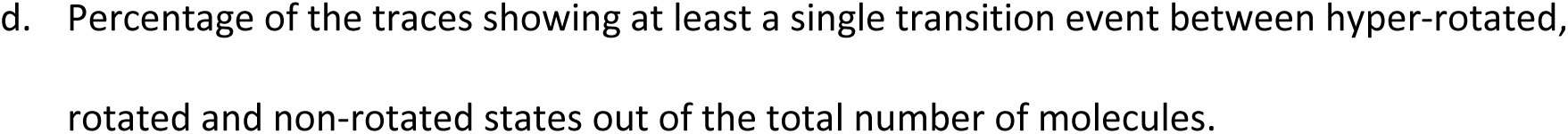
Statistical analysis for all tested ribosomal complexes.

Pre-translocation complexes (PREs) represent an intermediate step in translation where the A-site tRNA contains an elongated peptide chain while the P-site tRNA is deacylated. In our experiments, PREs were formed by adding N – Ac – Phe-tRNA^Phe^ non-enzymatically to the A-site with tRNA^fMet^ in the P site. The addition of an A-site tRNA had a modest effect on the conformational equilibrium and the percentage of dynamic molecules (Table 1). Nevertheless, the hyper-rotated state was significantly present (∼ 25%), with a concomitant increase in the non-rotated state population (Figure 4H, Table 1). Additionally, our data shows a slightly higher population of the rotated state as compared to previous results ^18^ (∼58%, previously 40%).

**Figure 4.**
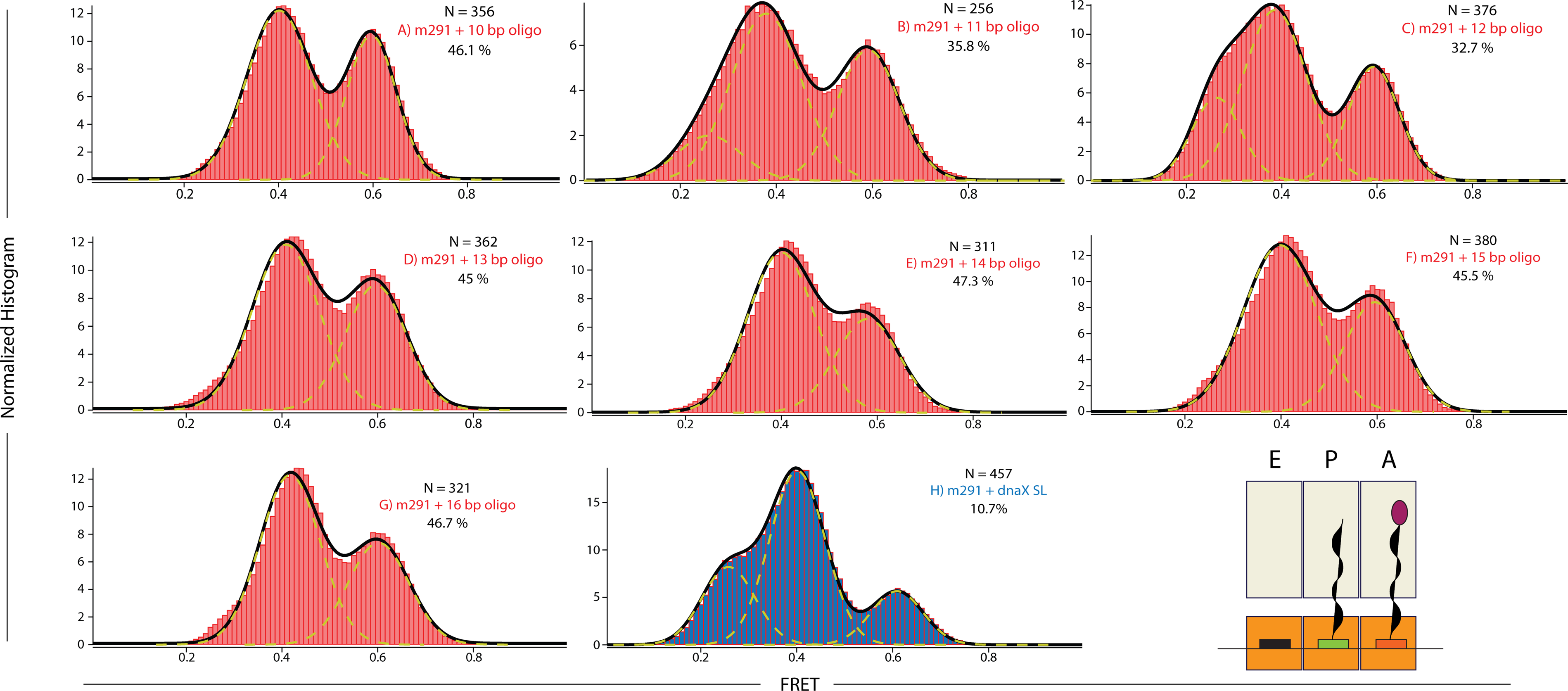
Normalized histograms complied from hundreds of traces showing distributions of FRET values for S6-Cy5/L9-Cy3 ribosomes containing a deacylated tRNA^fMet^ in the P site, N – Ac – Phe – tRNA^Phe^ in the A site and nucleic acid duplexes. N number of molecules used to compile each histogram; yellow lines represent individual Gaussian fits centered at ∼ 0.26 (hyper-rotated state), 0.38 (rotated state) and 0.58 (non-rotated state) FRET efficiency, black lines represent the sum of two (A, D-G) or three (B, C and H) Gaussians, % represent the percentage of fluctuating molecules for a given ribosomal complex.

**Figure 5.**
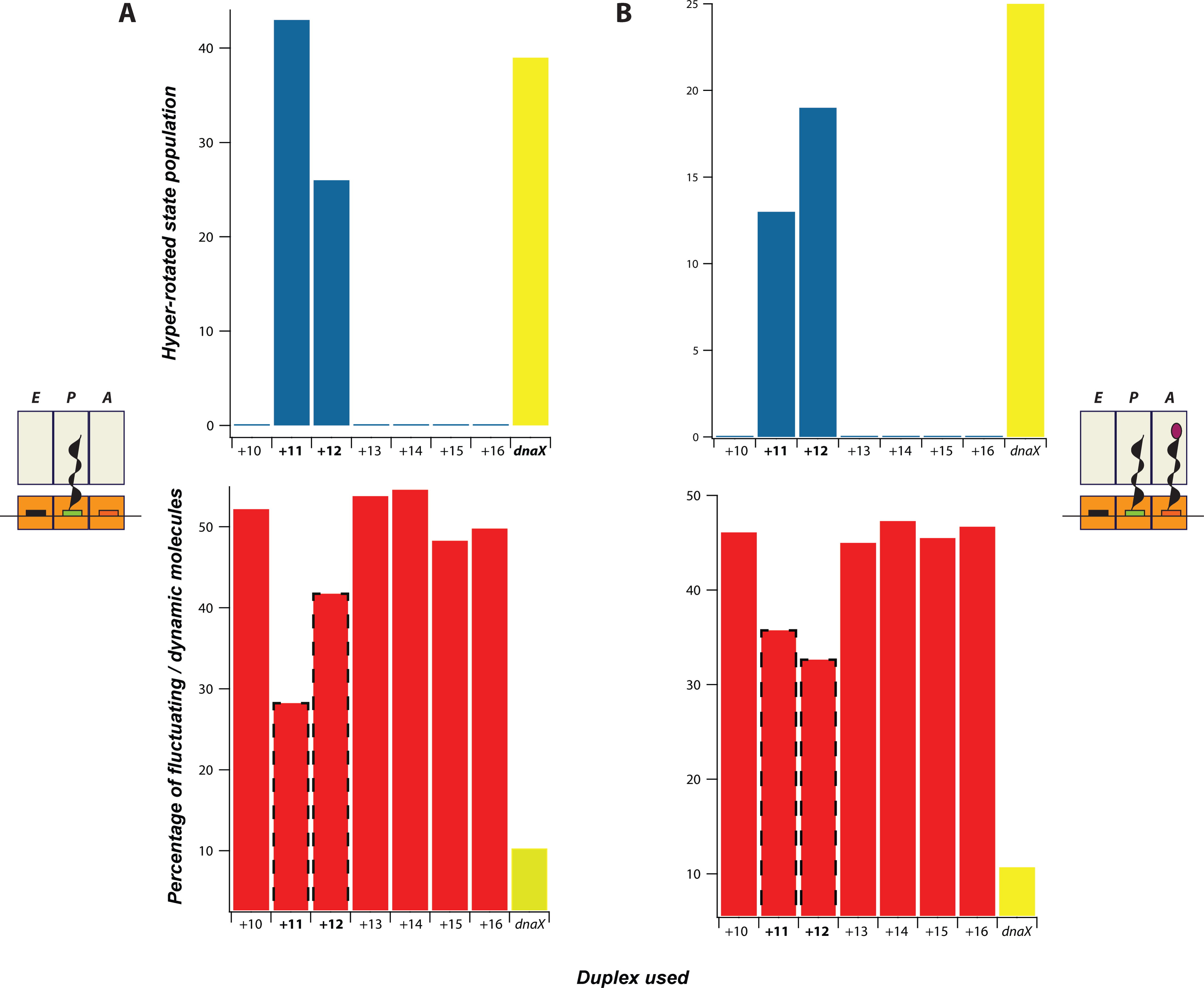
The negative correlation between the prevalence of the hyper-rotated population and the percentage of dynamic molecules. A) The left panel shows data from the singly-occupied P site, B) The right panel shows data for a pre-translocation complex. Red bars represent the percentage of fluctuating molecules. Blue bars represent the population of the hyper-rotated state. Highlighted data: *dnaX* is bolded and shown in yellow, +11 and +12 are bolded and highlighted with a dotted margin.

Likewise, the total percentage of dynamic molecules showed a significant drop as compared to the linear control but was comparable to the singly bound ribosomal complexes (∼ 10%). Our observations confirm the presence of a significant conformational change upon the presence of a structured hairpin in close proximity to the entrance tunnel. The data is comparable to previously published work and to the control, m291 ^17,18^.

### Duplex-walking and the induction of the hyper-rotated state

We used a duplex-walking assay to monitor the induction of the hyper-rotated state. A DNA – RNA helix was formed downstream of the ribosome binding region by hybridizing a complementary DNA oligomer to the mRNA (Figure 2). Shown previously *in vitro*, the helicase activity of the bacterial ribosome does not discriminate against RNA-RNA vs DNA-RNA duplexes; acting on both with a comparable unwinding efficiency ^9^. Seven DNA oligomers were designed to be of the same length, 20-mers, and have comparable stability (Table 2). The first DNA – RNA duplex was formed 10 nucleotides away from the +1 nucleotide in the P site. The added oligonucleotide was pre-hybridized to the mRNA during assembly and added in excess to the imaging buffer (1 µM, 1000X the concentration of the mRNA) (See Materials and Methods). Additional complexes were formed by walking different DNA-RNA duplexes one nucleotide at a time, in +1 increments, up to the +16 position (Figure 2B, Table 2). The chosen region, +10 to +16, covers the accessible region of the mRNA entrance tunnel based on X-ray crystallography data and spans the length of the tunnel to the outside rim ^31,32^.

**Table 2.**
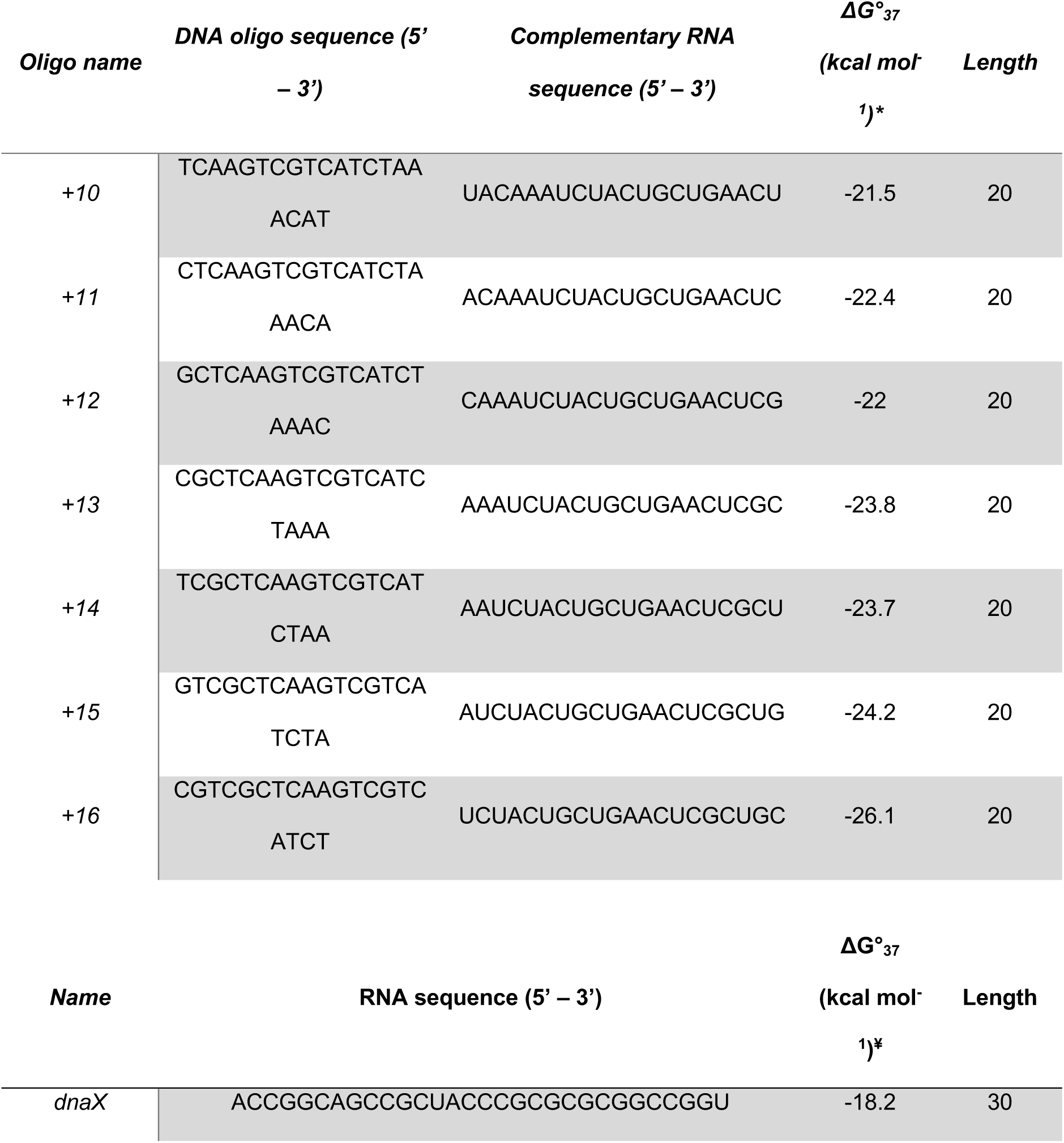
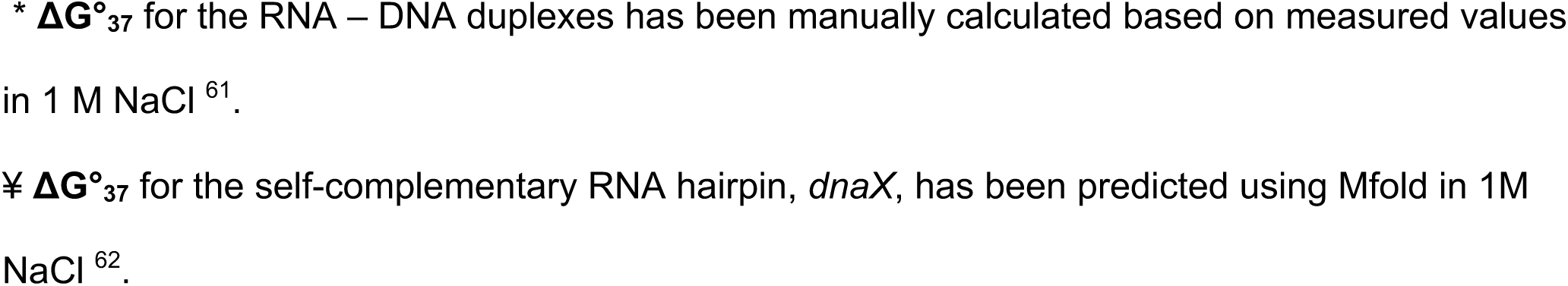
List of oligonucleotides used and their Gibbs free energy (kcal mol^-^. ^1^)

Following the same experimental design as for the *dnaX* hairpin, the first ribosomal complex was assembled with a single deacylated tRNA^fMet^ in the P site, and m291 pre-annealed to a complementary DNA oligo spanning the +10 to +29 region (Figure 2B). Binding of a single deacylated tRNA in the P site, drove the rotational equilibrium to favor the rotated conformation (∼70%) as per the linear unhybridized control (Figure 3A – H, Table 1). As shown previously, the assembled complexes show spontaneous fluctuations between the rotated and non-rotated conformations (∼52%) ^17^. Only when the DNA-RNA duplex was placed 11 or 12 nucleotides away from the P site, were we able to induce hyper-rotation (Figure 3B-C). When the +11 oligo was bound to the ribosomal complex, we observed a significant redistribution of the rotational equilibrium of the ribosome, where ∼ 43% of the ribosomal constructs were observed in the low FRET, hyper-rotated state versus ∼ 38% for the rotated state and ∼ 19% for the non-rotated state (Table 1). The appearance of the hyper-rotated state was concurrent with a drop in the percentage of fluctuating / dynamic ribosomes to ∼ 28%. Shifting the duplex downstream by a single nucleotide (+12 oligo), resulted in a reduction of the hyper-rotated population to ∼ 26% with a concomitant increase in the populations of the rotated (∼ 51%) and non-rotated (∼ 23%) conformations. The reduction in the hyper-rotated population was reflected in an increase in the percentage of fluctuating/dynamic molecules to 41.8%. Beyond the +12 position, +13 to + 16, except for few molecules sampling the hyper-rotated state, the low FRET, hyper-rotated population was suppressed to background levels and the return of the rotated and non-rotated conformations to their normal distributions. In the absence of the hyper-rotated state, the ribosomal constructs again displayed normal levels of fluctuation between the rotated and non-rotated states (∼52%), indicating DNA beyond the +12 position has little to no influence on intersubunit rotation.

Next, we wanted to investigate the influence of the A site occupancy on intersubunit rotation in the presence of a structured element. Generally, the addition of the A site tRNA shows a modest effect on the equilibrium distributions of the rotated vs non-rotated states, with a slight increase of the non-rotated conformation and a slight decrease in the percentage of fluctuating / dynamic molecules (∼ 45%) (Figure 4 A – H, Table 1) ^17^. To investigate any influences, a PRE was assembled by adding an N – Ac – Phe – tRNA^Phe^ to the A site. As shown above for single tRNA complexes, hyper-rotation was induced only in the presence of a DNA-RNA duplex placed 11 – 12 nucleotides away from the P site (Figure 2B). Surprisingly, the introduction of the A site tRNA resulted in a dramatic reduction in the observed hyper-rotated population in the presence of the +11 oligonucleotide (∼ 13%) as compared to the single P-site tRNA construct (∼ 43%) (Table 1). However, on the addition of the +12 oligonucleotide, we only observed a modest decrease (∼ 19%) in the hyper-rotated population as compared to its single tRNA counterpart (26%). As shown for the single tRNA data, the introduction of DNA oligonucleotides at the +13 position and beyond, up to +16, did not affect the equilibrium distribution between the rotated and non-rotated conformations, and we observed a suppression of the hyper-rotated state. These results show that a DNA – RNA duplex, spatially positioned at a specific distance from the entrance tunnel (11 – 12 nucleotides away), induces the hyper-rotated state formation regardless of the occupancy of the A – site.

### The kinetics of intersubunit rotation in the presence of a folded structure

FRET distribution histograms are useful for elucidating thermodynamic equilibrium distributions. In order to extract kinetic information pertaining to the dynamic heterogeneity of fluctuating ribosomes, smFRET traces showing fluctuations were analyzed using Hidden Markov Modelling (HMM), extracting idealized FRET trajectories (Figure 6A). Next, individual data sets were categorized into three populations, hyper-rotated, rotated, and non-rotated, based on a thresholding algorithm and cut-off limits (see Materials and Methods). Dwell times, time spent within a specific state before transitioning to a different state, were calculated. The first and last dwell times of each trace were discarded to minimize the bias, since the origin and/or the termination transitions were undetectable, respectively. Next, calculated dwell times were binned and plotted as a cumulative probability distribution. The data were then fitted to a single exponential. Kinetic rates were extracted from the fitting parameters (Figure 6B & 6C).

**Figure 6.**
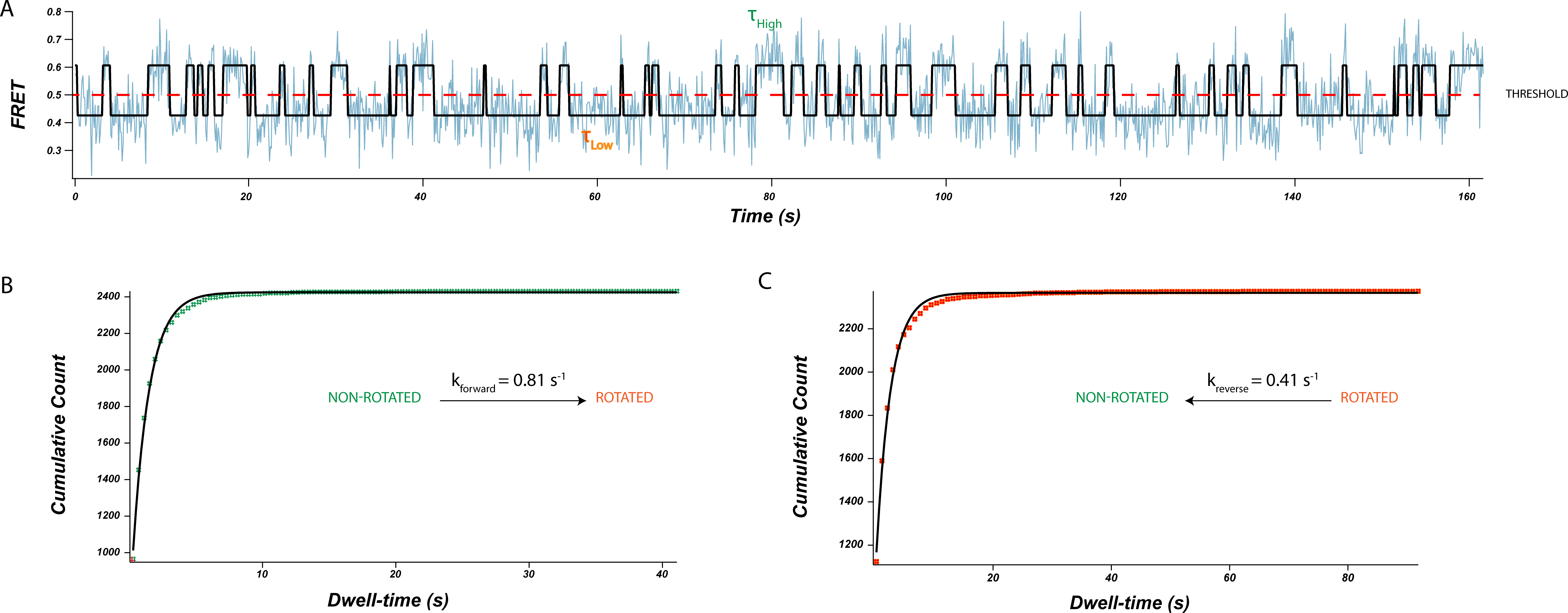
A) A sample FRET trajectory of a single-molecule with discrete FRET states identified via hidden markov modeling (HMM). Solid black line represents the hidden markov model fitting of the data. 𝝉 High represents the dwell time of the non-rotated state, 𝝉Low represents the dwell time of the rotated state. Dwell time is the time spent in a specific state before transitioning to a different state. B & C) Dwell time analysis after determining dwell times using a combined approach of HMM and thresholding. Measured dwell times from all traces were binned and plotted as a cumulative distribution and then fit with single-exponential functions using IgorPro. Kinetic rate constants for the transition from the non-rotated state to the rotated state (B), or the rotated state to the non-rotated state were calculated directly from the exponential fit (s-1).

Due to the limited observation window and the static nature of the hyper-rotated state, molecules transitioning from the rotated state into the hyper-rotated state showed only infrequent transitions, and thus were removed from our analysis. However, we did note the presence of these events in the collected traces (a total of 330 out of 5827 and 36 dwell times for all the ribosomal complexes with the P or the P and A site tRNA, respectively). The remaining fluctuating traces showed clear transitions between two distinct conformations: rotated and non-rotated states. Regardless of the presence or absence of nucleic acid duplexes downstream of the ribosome binding site, the measured kinetics of intersubunit rotation between the rotated and non-rotated states is unaffected (less than a factor of 2). The mean forward rate (Non-rotated ➔ Rotated) is 0.79 ± 0.06 s^-1^ and the mean reverse rate (Rotated ➔ Non-rotated) is 0.41 ± 0.1 s^-1^ at room temperature, under used *in vitro* conditions (Table 3). As compared to previously reported rates (NR ➔ R is 0.27 s^-1^ & R ➔ NR is 0.19 s^-1^), transition rates between the rotated and non-rotated conformations are higher which could be attributed to the higher activity of the ribosomes employed ^17^. Further investigation is required to detect forward and reverse transitions between the rotated and hyper-rotated conformations.

**Table 3.**
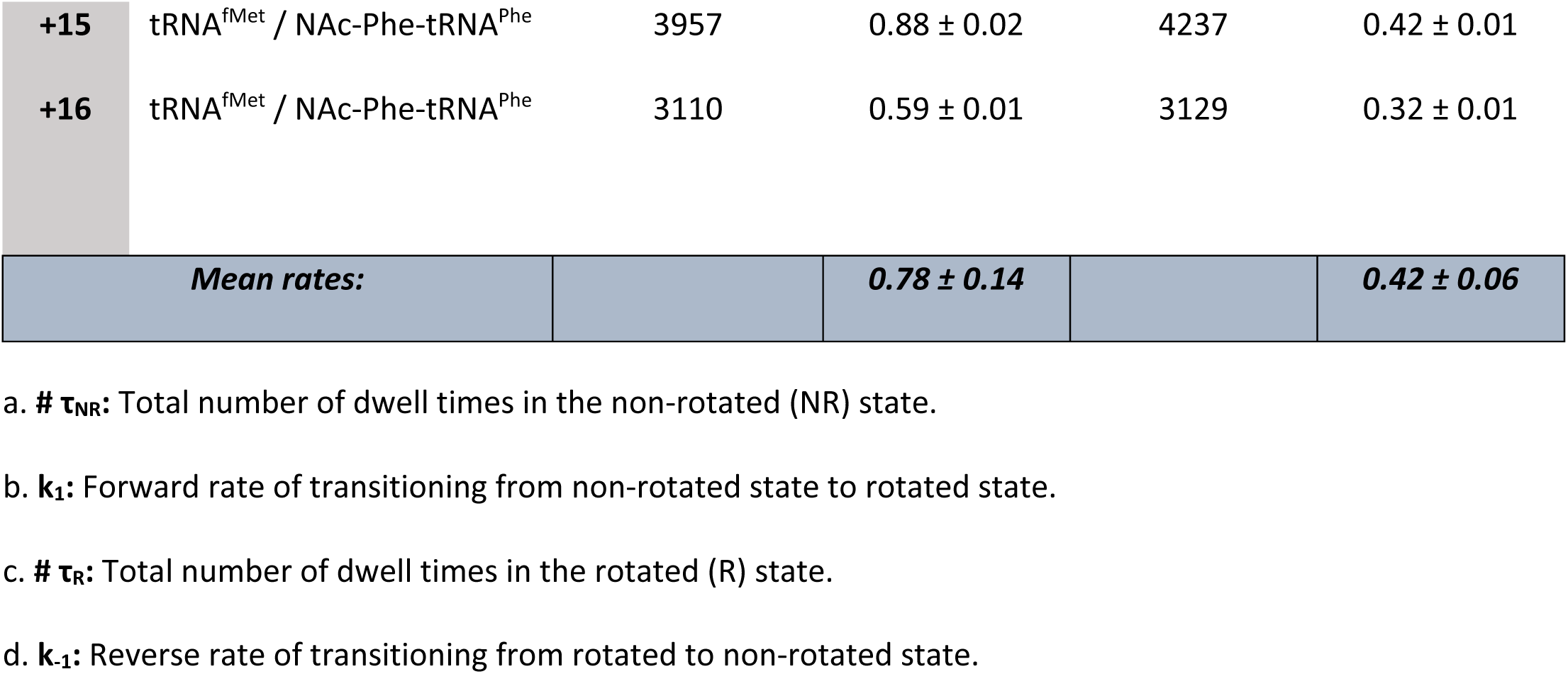
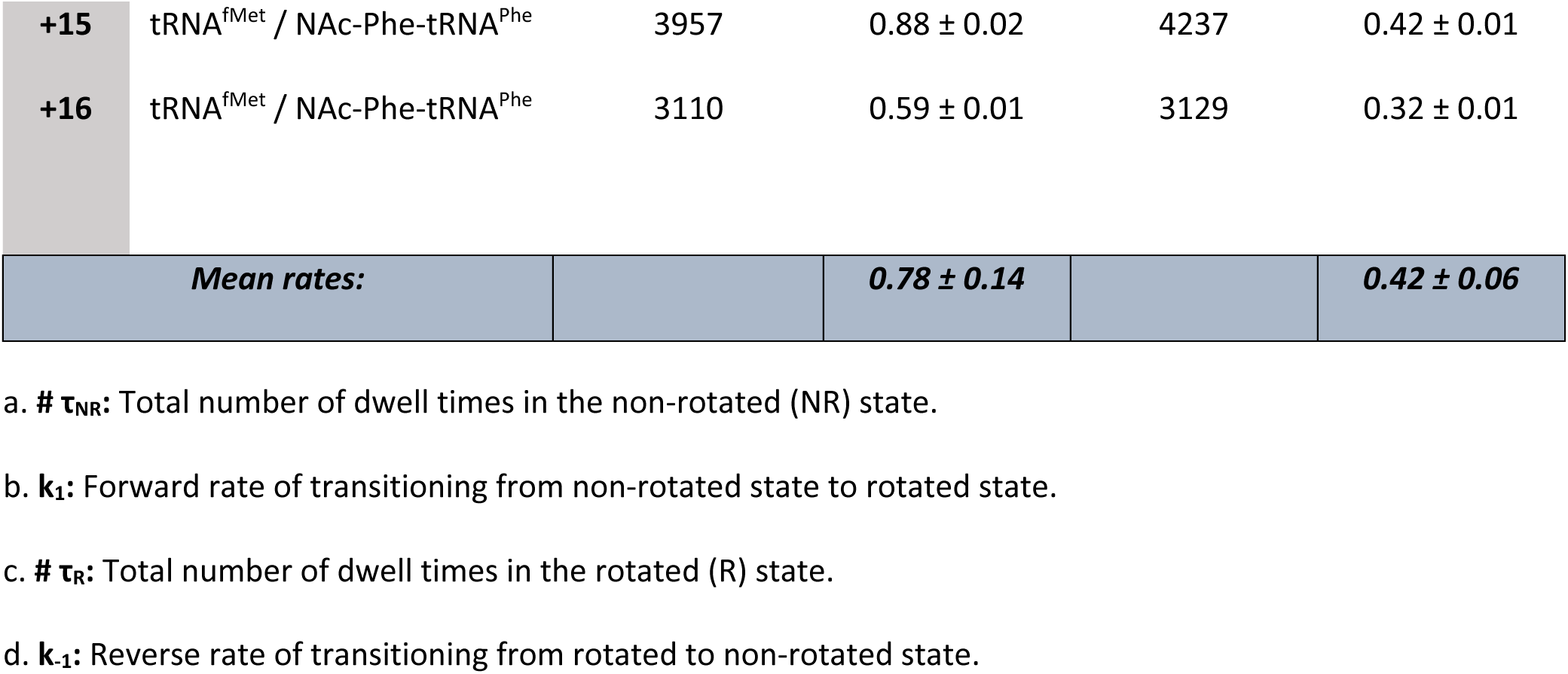
Kinetic rates measured for transitions between the non-rotated conformation and the rotated conformation.

### Molecular dynamics simulations show a change in ribosomal structure encountering structured mRNA

Molecular dynamics simulations were used to investigate how the proximity of structured mRNA to the entrance tunnel can affect the ribosomal structure and what could be the possible points of interaction between an incoming structured nucleic acid sequence and the mRNA entrance tunnel. Based on the structure 5AFI ^33^, we created three model systems of the *E. coli* ribosome that were similar to systems studied by Qin et al. (see Materials and Methods in Supporting Information for system preparation and MD protocol) ^18^. All systems contained the whole ribosome, including tRNAs at P and A positions, as well as the mRNA. Starting from +1 position to the final base at the position +30, the sequence of the mRNA in our model was identical to that in m291 (Figure 7A). The three systems differed by the position of the complementary DNA 15-mers, which were added to the mRNA strand at either +11, +13 or, +15 position (Figure S2). The systems are referred to as either +11, +13 or +15 system and three 100-ns-long simulations were performed for each system.

**Figure 7.**
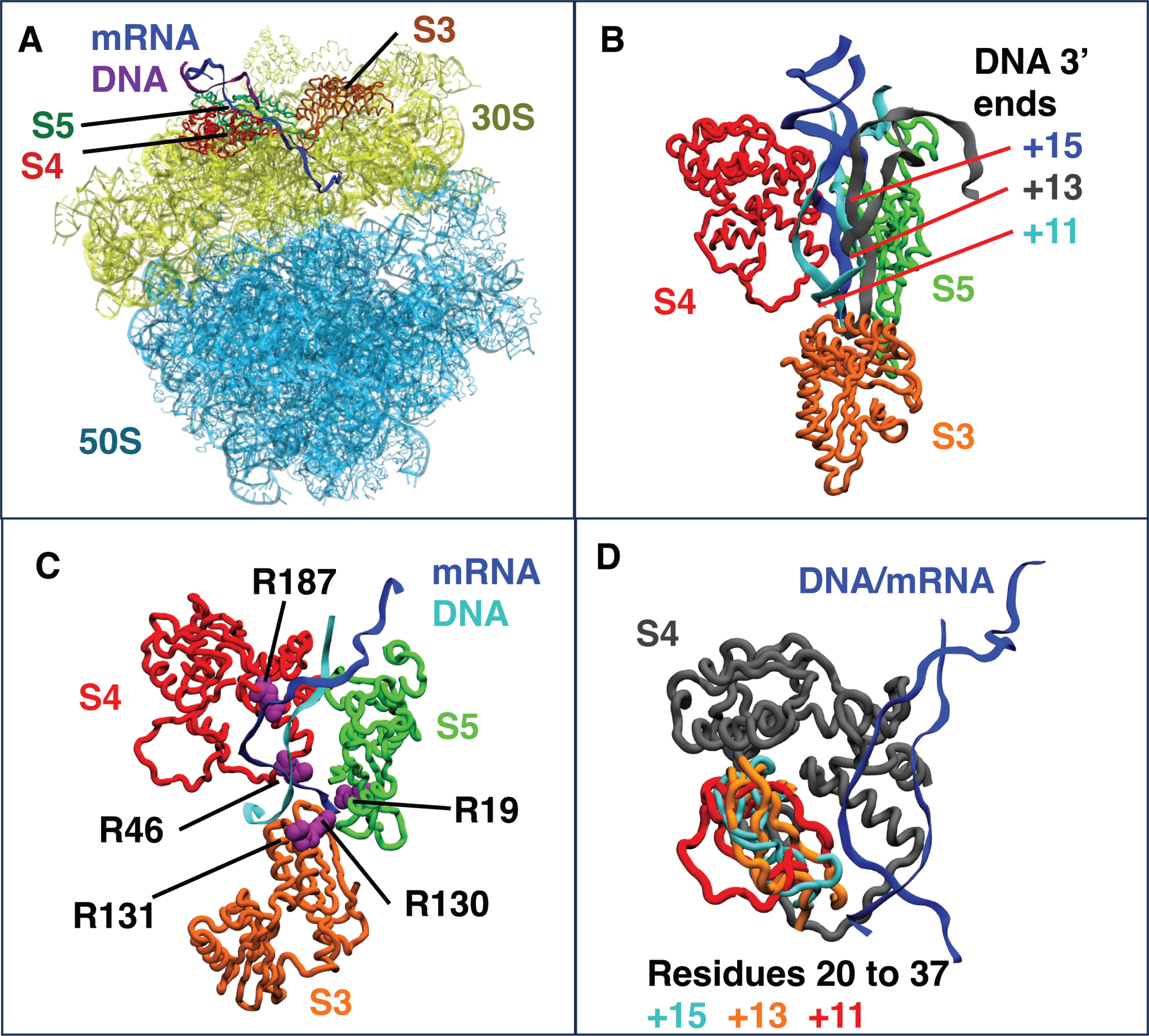
A) The ribosome model used in MD simulations with water and ions omitted for clarity. The 50S subunit is shown light blue and the 30S subunit is shown in yellow. The ribosomal proteins S3, S4, and S5 are colored orange, red and green, respectively. The mRNA is colored black, while the DNA 15-mer is colored blue. Proteins S6 and L9 used in the FRET experiments are shown in magenta and brown, respectively. B) The position of arginines in the ribosomal proteins S3, S4, and S5 that were found to maintain hydrogen bonds with either DNA or mRNA residues 50 to 70 in more than 50% of the trajectory frames for the +11 construct. The arginines are shown in cyan and, DNA and mRNA are shown in blue and black, respectively. Proteins S3, S4 and S5 are colored as in A.

All MD simulations showed high flexibility for the mRNA residues 50 to 70 and the complementary DNA strand. The DNA and mRNA strands remained in contact with each other for all three systems in all three simulations. However, the helical structure was not maintained by the end of any of the simulations of the +13 and +11 systems; in contrast, it was maintained in 2 out of 3 of the +15 system simulations (Figure S2), suggesting a higher structural stability of the latter. Although the DNA strand remained in the vicinity of S4 and S5 proteins in all systems, only the +11 system allows for interaction between the DNA strand and S3 (Figure S2).

Analysis of hydrogen bonding during the simulations found numerous interactions between DNA-mRNA strand and S3, S4, and S5 in the +11 system (Figure 7A, Table S2) that were maintained in more than 50% of the trajectory frames. Specifically, residues R130 from S3, R46 and R187 from S4, and R19 from S5 formed hydrogen bonds with mRNA residues 50 to 70 in the +11 system. R131 from S3 was the only hydrogen bond between the DNA strand and the ribosome. A significant reduction in hydrogen bonding between DNA-mRNA construct and S3, S4, and S5 proteins was found for +13 and +15 systems (Figure S3A). Hydrogen bonding with R130 and R131 from S3 and R187 from S4 were only transiently present in +13 and +15 systems (less than 35% of the frames), with R131 interacting with mRNA instead of DNA. Bonding with R46 from S4 maintains similar levels in +11 and +13 systems and is reduced in the +15 system. Finally, hydrogen bonding with R19 from S5 was absent in the +13 system and present at a similar rate in the +15 as in the +11. Previous work by Takyar et al. found that triple alanine mutation of R131, R132, and K135 in S3 and double mutation of R44 and R47 (our R46*) in S4 showed decreased helicase activity, whereas double mutation of R19 and R28 in S5 did not alter helicase activity ^9^. Our analysis suggests that residues R130 and R131 from S3, and R187 and R46 from S4 are likely to be important for helicase activity, based on persistence of these interactions in the +11 system and their significant reduction or complete absence in the less active +13 and +15 systems. The observed interactions with R19 from S5 appear less impactful because they are still present in the inactive +15 system to a similar degree as in the active +11 system. Additionally, analyzing changes in the structures of S3, S4, and S5, we noted significant changes in the fluctuations of S4 protein loop containing residues 25 to 35 in the +11 system (Figure S3B), suggesting that this loop could also be important for helicase activity. We also investigated structural changes in S6 and L9 proteins, which are labeled in the FRET experiments. The two proteins remained in the vicinity of each other in all of the +15 and +13 simulations, however a large separation of these proteins was observed for one of the +11 simulations (Figure S4). The distance between the FRET-labeled residues is not increased in this structural change, and significantly longer simulation times maybe be needed to observe the residue separation found in the hyper-rotated state.

## Discussion

Ribosomes undergo extensive conformational rearrangements during translation; the most notable of which is intersubunit rotation. Understanding these conformational changes accompanying translation and how the ribosome interacts with different elements, such as folded coding regions, provides a valuable insight into translation regulation mechanisms ^18,19,21,34^. Structured nucleic acids introduce a high energy kinetic barrier to the ribosome, thus pausing translation; a process exploited in co-translational folding, co-localization, and recoding mechanisms.

Inspired from work by the Noller lab, a DNA – RNA duplex was placed at variable distances from the ribosomal P site to determine the positional effect of the duplex on ribosomal conformation. We observed that structural elements close to the mRNA entrance tunnel drive the subunits into a hyper-rotated state, consistent with previous observation ^18^. The results show a stable low FRET state (∼ 0.22), as compared to the well characterized rotated (∼ 0.38) and non-rotated states (∼ 0.58). The determinant factor for the observed FRET states was decidedly the distance of the nucleic acid duplex to the ribosome. Under our experimental conditions, 7 to 8 nucleotides spacers from the P site were required to induce hyper-rotation. Such spacers place the nucleic acid duplex at the +11 to +12 position from the P site, a region close to the proposed helicase center of the ribosome. The +11 to +12 alignment is in agreement with the required spacer length for a folded mRNA structure to efficiently induce frameshifting ^35^. +13,+14, +15, and +16 spacers were ineffective in inducing a significant percentage of ribosomes in the hyper-rotated conformation. In addition, we observed a significant reduction in the hyper-rotated population for the +12 position but not for the +11 position. It is possible that this decline in the hyper-rotated population is due to the occupancy of the A site. Transitioning from initiation to elongation is accompanied by a conformational change of the shoulder of the 30S subunit which lays at an interface of the 30S and 50S subunits and near the mRNA entry tunnel. This domain closure leads to the contraction of the downstream tunnel and thus constricts its diameter by ∼ 1 – 3 Å ^32^. Although it remains to be determined, the tunnel narrowing accompanying the addition of N – Ac – Phe - tRNA^Phe^ to the A site may limit the accessibility of the oligo +12 from the P site.

Folded nucleic acid structures impose a kinetic and thermodynamic challenge to the ribosome. This challenge becomes particularly intriguing when ribosomes encounter stable downstream structured nucleic acid causing interaction of mRNA entrance tunnel proteins with the transcript ^16^. Recent smFRET studies have provided insight into the conformational dynamics of ribosomes during translation ^18,19,21,30,38^. For instance, ribosomes translating frameshifting signals with a downstream hairpin exhibit significant pausing events. These pauses have been linked to a non-canonical rotated conformation ^19^. Additionally, it has been demonstrated by Bustamante et al. through optical tweezers that the rate limiting step in structural interactions is not translocation, but rather pausing in a likely attempt to unwind mRNA interactions. Increasing the pulling force on either end of the mRNA, lowered the amount of pausing observed ^20^. In our investigation, we found that the hyper-rotated conformation represents a static state, while dynamic equilibrium exists between the rotated and non-rotated conformations. The kinetics of this pre-existing dynamic equilibrium remain unaffected even in constructs capable of inducing hyper-rotation. However, due to infrequent traces capturing the transition from the rotated to the hyper-rotated state, this shift occurs over a longer interval than the dynamicity expressed between the rotated and non-rotated states. We propose that this metastable local minimum in the ribosome’s energy landscape corresponds to the long-paused state, which has been previously observed by Puglisi et al. and referred to as a long-paused non-canonical rotated state ^19,21^. Unlike the studies conducted by Puglisi et al., which study active translation, we sought to precisely examine the effects of intersubunit rotation by utilizing stationary ribosomal complexes.

At equilibrium, and in the absence of translation factors, bacterial ribosomes rotate freely between the rotated and non-rotated states. Maintaining the delicate balance at equilibrium is crucial for translational fidelity. Furthermore, it has been demonstrated by Horan and Noller that translation is impossible when intersubunit rotation is restricted by the presence of a disulfide bond formed between small subunit protein S6 and large subunit protein L2. Binding of several translation factors and antibiotics (viomycin, sparsomycin, lincomycin, clindamycin, and chloramphenicol) at the A site preferentially stabilizes one conformation or the other ^17,26,36^.

Previously and within our current studies, the dynamic equilibrium between the rotated and non-rotated state in ribosomes encountering stable hairpins in close proximity of the mRNA entrance tunnel becomes disrupted, shifting a population into the hyper-rotated conformation ^18^. Seemingly, the close presence of structured elements at the mRNA entrance tunnel is sensed and transduced across the ribosomal network propagating a series of conformational changes resulting in hyper-rotation. Perturbations to the rotational freedom could disrupt the resulting allosteric communication network throughout the ribosome affecting the binding affinity of translation factors ^34,37,38^. The consequences of perturbation in rotational freedom may impact the fidelity of translation and frame-maintenance, a property that could be exploited in recoding mechanisms such as PRF. In other words, stably folded structures elevate the thermodynamic barrier such that a pool of molecules is shifted to the hyper-rotated conformation. Due to this thermodynamic barrier, we wanted to determine what interactions the ribosome would have with the nucleic acid structure.

Our simulation data showed structural differences in +11, +13, and +15 systems and interactional difference between the oligos and the ribosome, in agreement with previous results of Qin et al. ^18^. Notably, helical structure of the DNA-mRNA construct was only maintained in the +15 system, suggesting some degree of unwinding in the +13 and +11 systems. We also found that the DNA was sufficiently close interact to the S3 protein at the +11 position but not at the +13 and +15 positions, suggesting the importance of this protein for helicase activity. Specific interactions were observed between the mRNA-DNA construct and positively charged residues of the ribosomal proteins, which were most prominent in the +11 system (Table S2). We propose that the following residues could play a role in mRNA unwinding, allosteric communication, and subsequent hyper-rotation: R130, R131 from S3, and R46, R187 from S4. The S4 loop containing residues 25 to 35 could also play a role in the unwinding process, based on movement of these residues in the +11 system (Figure S3). Consistent with previous results from Takyar et al., the importance of residues R130, R 131 of S3 and R46 of S4 in ribosomal helicase activity have been shown experimentally^10^. However, the effect of S4 R187 has yet to be demonstrated. Thus, residues R130, R131 from S3, and R46, R187 from S4, then may serve as integral parts of the helicase center. While the studies carried out by Takyar and colleagues utilized mutations of S3 and S4, many other mutations occur on proteins in the helicase center resulting in the phenotype of ribosomal ambiguity ^9^. Ribosomal ambiguity mutations (ram) result in high rates of inaccuracies in the ribosome’s ability to incorporate the correct amino acid. Several mutations in S4 and S5, including D49Y and E68D in S4, have been shown to exhibit ribosomal ambiguity, and thus may have a larger capacity to enable hyper-rotation due to difficulties in unwinding downstream structures, something which would be interesting to experimentally verify ^39^. In our simulations we don’t observe any significant interactions between the mRNA/DNA construct and known ram mutations, suggesting that these mutations are more likely to alter S4 and S5 structures and the interface between the two proteins. These structural changes could in turn diminish the helicase functionality of the ribosome.

Recently, a cryo-EM structure has been released from Ermolenko et al. depicting the HIV frameshifting sequence hairpin bound to the A site of the 70S ribosome. In addition to this structure, S6/L9 labeled ribosomes were used to determine intersubunit rotation dependent of A site tRNA occupancy in both *dnaX* and HIV frameshifting sequences. While no hyper-rotation is reported, a few key experimental factors can explain this observed absence. As shown in our duplex-walking experiments, the contingency of the hyper-rotated state is dependent upon the distance of the nucleic acid duplex to the ribosome. Additionally, analysis of these data is challenging due to the hyper-rotated state exhibiting low FRET close to that of noise. Our results show that hyper-rotation occurs within the confines of an interaction between the helicase center of the ribosome and a frameshifting hairpin. As the A site is reported to be occupied by the HIV hairpin in ∼64%, of molecules, the possibility of observing hyper-rotation would greatly decrease. Indeed, it is observed that the HIV hairpin exhibits lower A site tRNA binding than that of the *dnaX* hairpin, a result which could arise from the competitive A site binding due to hairpin occupancy. Although a few structures of frameshifting hairpins bound to the A site exist ^12,40^, it remains unclear how the hairpin would move from outside of the ribosome to inside of the ribosomal A site during translation, and furthermore, how such a hairpin could move back outside the ribosome.

The hyper-rotated metastable state presumably lies within a deep energy well on the ribosome’s energy landscape with a high energy barrier (> kBT), in contrast to the lower energy barrier allowing the dynamic interconversion between the rotated and non-rotated states (∼ kBT)^41^. Thus, molecules trapped in this state are eliminated from the total pool of molecules in dynamic equilibrium as evidenced by a reduction in the percentage of dynamic / fluctuating molecules. We found a significant reduction of dynamic molecules when the self-complementary *dnaX* hairpin was utilized rather than a nucleic acid duplex with two fraying, and less stable, ends. Thus, structures that are more difficult to unwind, like *dnaX* hairpin, could possibly restructure the energy landscape, directing more molecules into the hyper-rotated state. As we show, even single-nucleotide displacements (+11 and +12 versus +13 and beyond) may significantly restructure the rugged energy landscape to divert ribosomes into different local pockets as evidenced by the appearance/disappearance of the hyper-rotate state. Similarly, EF – G -bound GDPNP and the antibiotic viomycin, respectively, are capable of structurally diverting the vast majority of ribosomes into the rotated conformation and raising the surrounding energy barrier (> kBT) enough to suppress the thermally driven intersubunit rotation ^17^. Identifying ribosome-mRNA interactions on residues of S3 and S4 through MD simulations leads us to believe that the interplay of mRNA-entry proteins with nucleic acid structure may have a significant impact on hyper-rotation.

Despite the mounting knowledge of the intrinsic helicase activity of the ribosome and the conformational changes accompanying translation, many of the mechanistic details are missing. The intricacies of local structural rearrangement around the mRNA entrance tunnel are essential in how ribosomes sense and unwind structured mRNA elements. How, then, would the ribosome transduce such signals allosterically throughout the macromolecular complex? Further investigations are required to address the remaining challenges and questions involving the impact of the stability of structured nucleic acids on the conformational dynamics of the ribosome.

## Materials and Methods

### Ribosome preparation

We introduced specific cysteine mutations via site-directed mutagenesis to recombinant ribosomal proteins S6 (D41C) and L9 (N11C). Next, we conjugated fluorescent reporters Cy3 and Cy5 maleimide to S6 (D41C – Cy5) and L9 (N11C – Cy3).

Fluorescently labeled proteins were then reconstituted into ΔS6 / ΔL9 dual knockout His-tagged 70S Escherichia coli ribosomes ^30^. Testing the biochemical activity of the generated ribosomes using filter binding assays and puromycin-reactivity assays showed that the partially reconstituted ribosomes have comparable tRNA binding and EF-G translocation efficiency to wild-type ribosomes ^30,42^.

### Preparation of charged tRNA and mRNA

m291 and *dnaX* were transcribed using T7 RNA polymerase and purified on a denaturing polyacrylamide gel ^18,43,44^. m291 is a derivative of T4 bacteriophage gene 32 ^9^. *dnaX* hairpin is a derivative of the *dnaX* gene encoding DNA polymerase in *Escherichia coli* and, along with a slippery site and a Shine-Dalgarno sequence upstream, is involved in -1 programmed frameshifting ^8^. tRNA^fMet^ and tRNA^Phe^ were purchased from MP Biomedicals and tRNA probes, respectively. N – Ac – Phe – tRNA^Phe^ were aminoacylated using DEAE-purified S100 enzymes ^45–47^. Aminoacylated tRNAs were purified from the charging reaction and the extent of aminoacylation was verified by acid gel electrophoresis. DNA oligonucleotide variants were ordered from Integrated DNA Technologies (IDT) (Table 3).

### Preparation of ribosomal complexes for smFRET

All ribosomal complexes were assembled in polyamine buffer (20 mM Hepes, pH 7.5, 6 mM MgCl2, 150 mM NH4Cl, 6 mM ßME, 2 mM spermidine, 0.1 mM spermine) ^17^. m291 was pre-annealed to the desired complementary DNA oligonucleotide and the biotin-labeled DNA oligonucleotide (5’ biotin – CTTTATCTTCAGAAGAAAAACC-3’, IDT) at 65°C for 5 min and then cooled on ice for 10 min. To initiate ribosomes with a P-site tRNA only, 1 μM S6/L9 labeled ribosomes were incubated with 2 μM of the pre-annealed m291 / DNA duplex and 2 μM tRNA^fMet^ at 37 °C for 20 min. Pre-translocation complexes were prepared in a similar manner, except including incubation of the A site tRNA, N – Ac – Phe – tRNA^Phe^ at 37°C for an additional 20 min. *dnaX*-containing samples were prepared following the same protocol with the elimination of the duplex forming DNA oligonucleotide in the pre-incubation step.

Quartz slides were pre-cleaned and passivated with a mixture of m – PEG – succinimidyl valerate (MW 5000) and Biotin – PEG – succinimidyl valerate (MW 5000) (Laysan Bio Inc.) and pretreated with neutravidin (0.2 mg/mL) (Pierce) ^18,48,49^. Ribosomal complexes were diluted to a final concentration of 1 nM and immobilized on a quartz slide (**Error! Reference source not found.** A). To minimize photobleaching, all samples were imaged in polyamine buffer supplemented with an oxygen-scavenging system; 0.8 mg/mL glucose oxidase (Sigma), 0.625% β-d-glucose, 0.02 mg/mL catalase (Roche), and 1.5 mM Trolox (Sigma) ^17,48^. Duplex-forming DNA oligonucleotides complementary to the desired region were added to the imaging buffer to a final concentration of 1 μM.

### Acquisition and analysis of smFRET data

A 532 nm laser (Spectra-Physics) was used to excite the donor dye (Cy3) via prism-based total internal reflection microscopy (TIRFM) ^48,49^. The resulting fluorescence emission was split into two pathways (Cy3 and Cy5 emission) with a 630dcxr dichroic mirror (Chroma). The emission signal was collected via an Andor iXonEM + 897 Electron Multiplying Charge Coupled Device (EMCCD) camera. Movies were recorded using Single (custom software generously provided by Taekjib Ha) at an acquisition rate of 100 ms. A calibration image was acquired using crimson fluorescent FluoSpheres® (Invitrogen). The movies were processed to extract donor and acceptor intensities using custom IDL scripts.

Next, data was sorted and analyzed using custom Matlab scripts (Mathworks). The acquired traces were sorted and validated based on predefined criteria: only traces showing single step photobleaching for both dyes, Cy3 and Cy5, were considered; the trace had a minimum of 10 data points (1 s). Traces showing positive correlation between the intensities of the donor and acceptor dyes, intensities less than 100 arbitrary units (au) or FRET states less than a set threshold (0.15) were manually excluded. Each trace was normalized to its temporal length to ensure equal contribution to the final histogram ^18,50^. Data points were smoothed with a five-point window average. A comparable number of molecules were selected to contribute to the histogram. Peaks within the normalized histograms were identified automatically via the peak finder function of IGOR Pro (Wavemetrics) and fitted to either two or three Gaussian distributions ^18^. The viability of the fit was partially assessed from the consistency of the width of the individual peaks.

For rate determination, a combined approach was used. Discrete FRET states were identified within traces using Hidden Markov Modelling (HMM, vbFRET) ^51^. Next, dwell times of the different states were categorized into three bins (hyper-rotated, rotated and non-rotated) based on a thresholding algorithm (custom Matlab script) ^52^. Individual dwell times were then calculated and plotted as a cumulative probability distribution that was then fit to a single exponential curve using IGOR Pro (Wavemetrics).

### Molecular Dynamics Simulations

To prepare the models of the ribosome, we started with the structure of the *E. coli* ribosome (PDB code 5AFI) that contains three tRNAs at the A/T, P, and E positions ^33^. The tRNA in the E site was deleted to match the ribosomes used experimentally. Because the mRNA strand in this structure was very short, we also used the PDB structure 4V4Y that contains a full mRNA strand in the *Thermus thermophilus* ribosome ^53^. We aligned the 16S segments in 5AFI and 4V4Y, which have very similar structures, to obtain the correct positions of the mRNA strand and the Shine-Dalgarno region of the 16S for our model. The 16S rRNA and the mRNA in 5AFI were replaced with the ones from 4V4Y after alignment to 16S. The replaced structures were minimized in the *E. coli* ribosome to eliminate any steric clashes. We used the cIonize plugin ^54^ in VMD to neutralize the system by adding Mg^2+^ ions with completed octahedral solvation shells at points of electrostatic potential minima. The last base in the mRNA strand was at the +30 position; between positions +1 and +30, the bases were mutated to the ones in the 291mRNA. We added the DNA 15-mers by first using the Nucleic Acid Builder ^55^ to generate the structures of 291mRNA starting from the +13 dposition and the complimentary DNA 15-mers at either +11, +13, or +15 position. The three generated structures were equilibrated in water for 10 ns, after which they were covalently linked to the mRNA in our *E. coli* ribosome system at the corresponding position, resulting in the +11, +13, and +15 systems. Bulk water was added with the VMD Solvate plugin, after which chloride and potassium ions were added at a concentration of 0.1 mol/L with the Autoionize plugin.

All molecular dynamics (MD) simulations were performed with NAMD ^56^. The CHARMM36 protein ^57^ and nucleic acid ^58^ force fields were used for the ribosome, whereas the TIP3P model was used for water ^59^. We used the particle-mesh Ewald method to calculate long-range electrostatics ^60^. A cutoff of 12 Å was used for the van der Waals interactions and a potential switching function was applied from 10 to 12 Å to ensure a smooth decay to zero. All covalent hydrogen bonds were kept rigid, which allowed us to integrate the equations of motion with a 2-fs time step. The temperature and pressure were kept constant at the biologically relevant values of 310 K and 1 bar with a Langevin thermostat and piston, respectively. For hydrogen bond analysis, bonds were defined as present if the acceptor-donor distance was 3.5 Å or less and bond angle was between 145° and 180°.

The constructed systems were minimized in two steps. In the first step, only water and ions were minimized, followed by minimization of all atoms in system in the second step. The equilibration was done in three 2-ns-long steps. In the first step, water and ions were equilibrated, after which mRNA bases in positions +1 to +3 and the complementary bases in the P-site tRNA were equilibrated, followed by equilibration of all side chains with only the backbone restrained in the final equilibration step. After equilibration, all systems were sampled for 100 ns in triplicate.

Table S1. Fraction of trajectory frames with hydrogen bonds present between the selected ribosomal protein residues and the DNA-mRNA construct. Errors are calculated from standard deviation between the three runs.

**Figure S1.** Representative trace of S6-Cy5/L9-Cy3 ribosomes containing a deacylated tRNA^fMet^ in the P site. A) Hyper-rotated state. B) Rotated state. C) Non-rotated state. D) A dynamic molecule fluctuating between rotated and non-rotated states. Cy3 and Cy5 intensities are shown in green and red, respectively. The calculated FRET curve is shown in blue. Solid black lines highlight the different FRET states identified in the trace using Hidden Markov Modelling (HMM).

**Figure S2.** Selected snapshots from the end of MD simulations comparing the structures of different constructs, A) +15 system. B) +13 system. C) +11 system. In all subfigures, the mRNA is colored black, while the DNA 15-mer is colored blue. The ribosomal proteins S3, S4, and S5 are colored orange, red and green, respectively. The locations of the DNA strands’ 3’ ends are marked with a magenta circle.

**Figure S3.** A) Root mean square fluctuations (RMSF) for the S4 residues in the +15, +13, and +11 systems. The black line shows the average between the three runs, and the red area shows standard deviation. B). Structure of S4 (red) with the loop of flexible residues 20-30 shown in gray. The position of DNA/mRNA (blue and black) relative to S4 is also shown.

**Figure S4.** Snapshots of S6 (magenta), and L9 (brown) from our simulation, the residues involved in the FRET experiments are shown in blue. A) A snapshot from +15 system representing a typical orientation of S6 and L9 in our simulations. B) Separation of S6 and L9 observed in one of our simulations for the +11 system.

